# Excitation energy transfer between higher excited states of photosynthetic pigments: 2. Chlorophyll *b* is a B band energy trap

**DOI:** 10.1101/2023.01.26.525641

**Authors:** Jan P. Götze, Heiko Lokstein

## Abstract

Chlorophylls (Chls) are known for fast, sub-picosecond internal conversion (IC) from ultraviolet/blue absorbing (“B” or “Soret” states) to the energetically lower, red light-absorbing Q states. Consequently, excitation energy transfer (EET) in photosynthetic pigment-protein complexes involving the B states has so far not been considered. We present, for the first time, a theoretical framework for the existence of B-B EET in tightly coupled Chl aggregates, such as photosynthetic pigment-protein complexes. We show that according to a simple Förster resonance energy transport (FRET) scheme, unmodulated B-B EET likely poses an existential threat, in particular the photochemical reaction centers (RCs). This insight leads to so-far undescribed roles for carotenoids (Crts, cf. previous article in this series) and Chl *b* (this article) of possibly primary importance.

It is demonstrated how pigments in a photosynthetic antenna pigment-protein complex (CP29) undergo FRET. Here, the focus is on the role of Chl *b* for EET in the Q and B bands. Further, the initial excited pigment distribution in the B band is computed for relevant solar irradiation and wavelength-centered laser pulses. It is found that both accessory pigment classes compete efficiently with Chl *a* absorption in the B band, leaving only 40% of B band excitations for Chl *a*. B state population is preferentially relocated to Chl *b* after excitation of any Chls, due to a near-perfect match of Chl *b* B band absorption with Chl *a* B state emission spectra. This results in an efficient depletion of the Chl *a* population (0.66 per IC/EET step, as compared to 0.21 in a Chl *a*-only system). Since Chl *b* only occurs in the peripheral antenna complexes, and RCs contain only Chl *a*, this would automatically trap potentially dangerous B state population distantly from the RCs.

## Introduction

Photosynthetic organisms have to balance between vastly differing irradiation conditions. Light is required for survival, yet photodamage may also be eventually lethal.^1–4^ Damage from too much light occurs mainly at the photosynthetic reaction centers (RCs).^5–11^ It has been shown that plants can cope with all of these challenges and are basically not limited by light energy input.^12–14^ A key factor in photoprotection is non-photochemical quenching (NPQ), often seen to be associated with the accessory chromophores (see below).^15–17^

### Photosynthetic pigments

In plants, the main pigments are chlorophylls (Chls) *a* and *b*. In photosynthetic bacteria, bacteriochlorophylls (BChls) are found. (B)Chls act as light harvesters and electron transfer chain components.^18,19^ (B)Chl absorption spectra in the ultraviolet/visible/near-infrared (UV/Vis/NIR) regions are shown in Figure 1A, together with solar irradiation spectra.^20^ Two main absorption bands are observed, called B (or Soret) and Q.^21^ Light absorbed by the “red” Q band is directly used for the two light reactions in the photosystems I and II (PSI and PSII).^8,9,22^ B band excitations are converted to lower energy Q excitations via internal conversion (IC). B → Q IC has been observed in Chl *a* to occur within approximately 100 to 250 fs.^23–25^ The Q and B bands both consist of multiple electronic states.^26–29^

**Figure 1:**
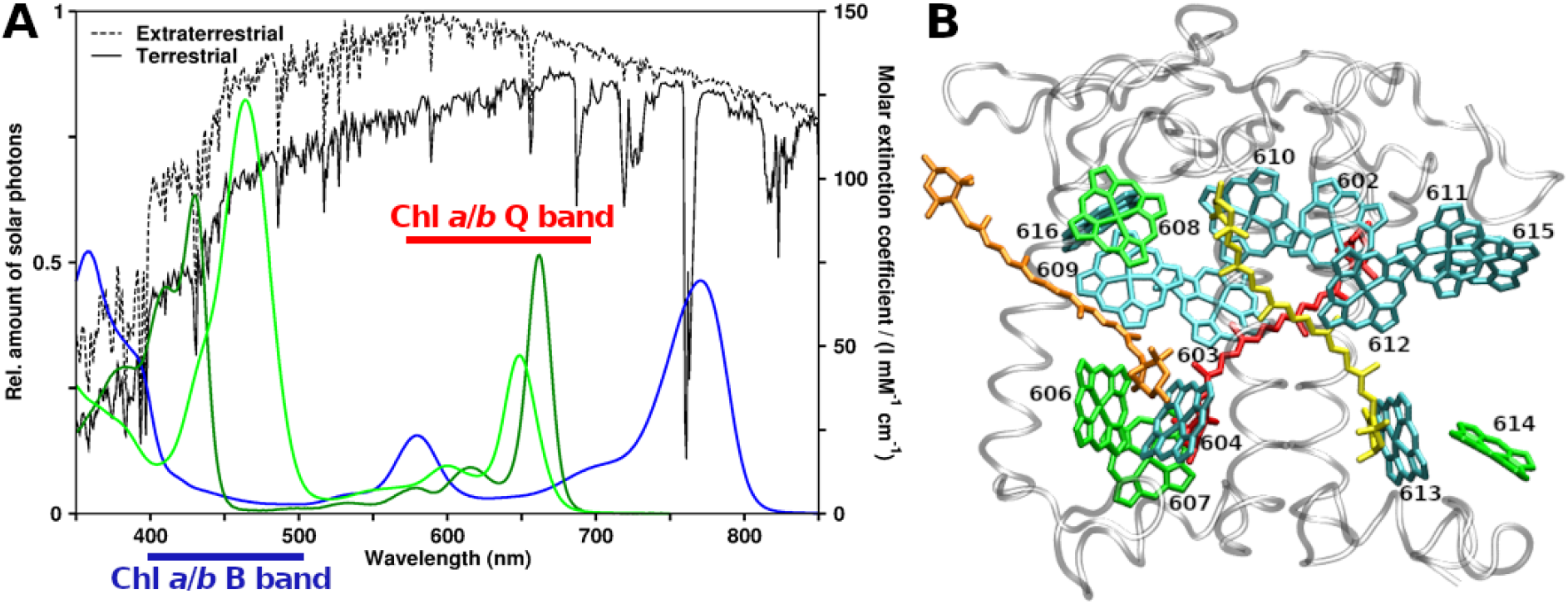
Spectroscopy and structural arrangement of typical plant photosynthetic pigments. A: Ultraviolet/visible region absorption spectra of Chls a, b and BChl a (dark green, light green and blue, respectively), compared to solar irradiation (from Gueymard 2004). B: CP29 antenna complex from Pisum sativum, PDB structure 5XNL by Su et al. (2017). Viewpoint along Thylakoid membrane plane, looking towards the center of the photosystem II supercomplex. Stromal side at the top, lumenal side at the bottom. Protein grey and transparent, Chls a shown as cyan, Chls b as green (Chl numbering as given in earlier CP29 structures, e.g., 3PL9). Crts also shown, lutein in yellow, neoxanthin in orange and violaxanthin in red.

Excitation energy transfer (EET) between (B)Chls may occur at short intermolecular distances in the form of coherent EET (electronic coupling) or, especially at longer distances, via incoherent EET. The underlying coupling can be modeled on the basis of dipole-dipole interactions by Förster resonance energy transfer (FRET) using a point dipole approximation (PDA).^30–35^ For the B band, there have been very few efforts so far to elucidate EET processes (see supporting information, SI). However, even assuming ultra-fast IC, coherent and incoherent EET have been shown to occur among B states of Zn-porphyrins (analogues of Chls), see SI.^36–38^

Neglecting B band EET for Chls relies on the assumption that Chl B → Q IC outpaces all other processes. The presence of carotenoid (Crt) absorption in the same spectral region poses an additional experimental obstacle. There have, however, been some experimental and theoretical studies which we will discuss shortly below.

### B band-EET in Chl assemblies

Zheng and coworkers^24^ simulated the excited state kinetics in a Chl *a* dimer,^39^ focusing on B → Q IC kinetics. A deexcitation pathway from B via Qx to Qy (the two electronic states forming the Q band) was proposed.

The simulation was started with a 50% population in an energetically higher B_high_ state, and 50% in a lower B_low_ state. From their simulations, it can be seen that the initial process is not B → Qx or B → Qy IC. Instead, a near-complete depopulation of the B_high_ state is found first (Figure 3 in Zhang *et al*.^24^). After about 10 fs, the initial 0.5/0.5 B_high_/B_low_ population distribution has reformed to 0.17/0.75 (the remaining 0.08 population being transferred to lower Q states). Thus, B → B EET outweighs IC from the B states into the Q band by 0.33/0.08 (population transferred/lost). Furthermore, 10 fs is already longer than the B → B EET lifetime in their simulations. This is, to our knowledge, the first observation of the efficacy of (coherent) B band EET, although the authors apparently did not note it as such.

Their study predicts ~100 fs for B → Q IC, in agreement with the experiment.^24^ All these findings strongly suggest that EET between B states precedes B → Q IC also in Chls, not only in Zn-porphyrins.^36–38^ From an open-minded physicochemical perspective, this is not surprising;^40^ FRET does not *a priori* exclude EET between states of higher energy.

Experimentally, B → B EET has also been found, although not receiving proper resonance in the literature. *Stepwise* two-photon excited fluorescence (TPEF) spectra^41,42^ demonstrate fluorescence signals from the B band. For Chl *a* and *b* in diethyl ether, the authors find B emission maxima at 440 and 462 nm, respectively. The authors further *observe* EET in the B band in light-harvesting complex II (LHCII), attributed to adjacent Chls *a* and *b*. In CP29, they find that the B band emission occurs primarily from the originally excited Chls. Thus, B band EET is apparently possible and system dependent. Recently, it was also (computationally) proposed for CP29 that the coupling pattern between Chls is state-dependent and steered by the protein environment.^43^

In the previous article in this series, these insights were put to the test, and it was shown that B-B EET appears to be a significant process in a CP29 FRET model. The main result was that B-B EET is strongly affected by the presence of Crts. The same analysis however indicated that Chl *b* might play an as of now unknown role for the B-B EET, beyond simply enhancing the spectral width of the overall Chl absorbance, *i.e*., a function beyond enhancing light harvesting. This article will analyze the effect of Chl *b* on FRET in a CP29 model.^44^

First, FRET of mixed Chl-Chl B band interactions for the isolated (non-CP29) chromophores is assessed. Then, the effect of Chl *b* on these processes in CP29 will be computed and combined with the effect of Crts. As a last step, the initial B band absorption competition between the CP29 pigments will be computed, before ending the article with a summary of the results.

## Methodology

### FRET model

We employ FRET theory, effectively only including dipole-dipole couplings.^40,45^ The total FRET efficiency for a donor (D) transferring to *i* acceptors (A) is

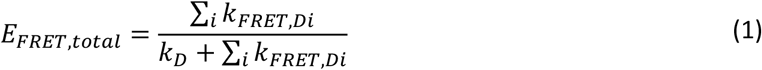

*k_D_* as the decay rate of the donor state in the absence of A (equal to 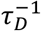, *τ_D_* being the state lifetime of the isolated donor, Chls *a/b:* 100/58 fs for B,^25^ 6.3/3.2 for Q,^46^ Crt bright S_2_ state: 163 fs^47^). Note that *k_D_* was reported to be nearly independent regardless of Chl *a* monomers or dimers (below 10% difference^24^), so extracting *k_D_* from the sum appears justified. *k_FRET,DA_* is the rate of EET *via* FRET

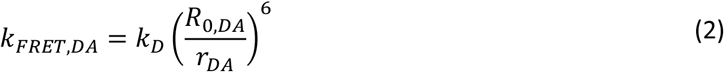

with *r_DA_* as the D-A distance and *R*_0,*DA*_ as the Förster radius. Defining *r_DA_* is not trivial for extended chromophores such as Crts and FRET can only be expected to provide qualitative answers for small distances.^48^ *R*_0,*DA*_^49^ is

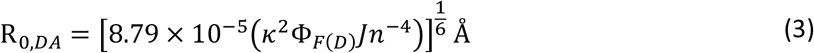

with Φ_*F*(*D*)_ being the donor quantum yield, *n* the refractive index of the medium (1.4 in a protein environment^50^), *J* the spectral overlap (in M^-1^ cm^-1^ nm^4^), and the orientation factor *κ* being

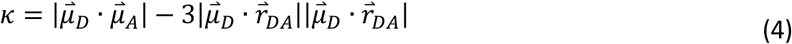

with 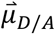 as the transition dipole moment (TDM) of the donor or acceptor state. For *κ*, random orientation corresponds to a *κ*^2^ of 2/3, while perfect arrangement is 4,^45^ to not bias our analysis, we will take both into account. TDMs for Chls were located along atomic axes, mimicking the results of the TDM orientations by Sirohiwal *et al*.^27^ for Chls (Q: along NB-ND axis, B: along C4A-C4C axis). For the bright state of Crts, S_2_,we relied on our earlier calculations,^29^ resulting in a TDM orientation along the axis connecting atoms C12 and C32. More elaborate approaches are possible, but not required for the proof of concept here.^51^ Molecular centers were defined for Chls to be the Mg ion position in chain R of the 5XNL cryo-EM structure.^44^ For Crts, molecular centers were computed as the averaged position of all conjugated carbon atoms of the Crt in question.

For the B band (and brightly absorbing Crt S_2_ state, see previous article), obtaining Φ_*F*(*D*)_ poses a challenge, and experimental estimates only present an upper limit of 10^-4^ for the Chls.^41^ For Crts, only a single value of 1.5 10^-4^ was found.^47^ In the previous article, a theoretical analysis^52,53^ provided slightly higher values, except for Chl *b*, for which a *Φ*_*F*(*Chl, b, B*)_ of 0.92 10^-4^ was computed. To bias against the processes investigated here, the lower values, as listed in Table S1 of the SI were chosen, just as in the previous article. This leaves the task to find a value for *J*, which is

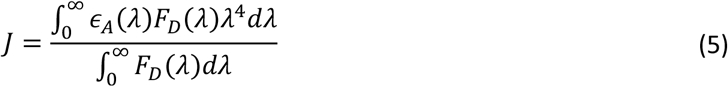

with *ϵ_A_*(*λ*) as molar extinction coefficient at wavelength *λ* of the acceptor, and likewise *F_D_* as the fluorescence of the donor.

For the Chl spectra, the emission spectra from Leupold *et al*. (B band)^41^ and Du (Q band)^54^ were used and the absorptions from Du (both bands).^54^ Q and B bands were separated at 470.5 nm for Chl *a* and 495 nm for Chl *b*. For the Crts, the same spectra were used as for the previous article of this series, from various sources.^16,29,55–58^

### Structural model

The structural model employed throughout the article is the CP29 peripheral antenna complex, using chain R of PDB entry 5XNL, a cryo-EM structure of PSII.^44^ All Chl and Crt sites are included, assuming a wildtype pigment composition first, followed by modifications of Chl *b* content and/or Crt presence. The corresponding site energies were taken from our earlier work,^43,59^ the values explicitly listed in the SI, Table S2. For Crts, no site energy shifts were applied. Intermolecular distances were used as listed in Tables S3 and S4 of the SI.

### Relative absorption in chromophore mixtures

To assess the competition between the different chromophores in CP29, the absorbed photon intensity *I*(*λ*) for a given irradiation spectrum *I*_0_(*λ*) is required for each chromophore in the system, applying the Lambert-Beer^60,61^ law

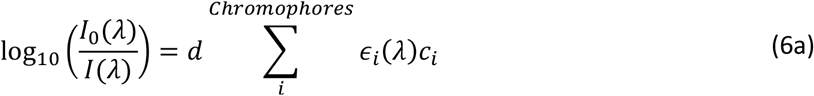

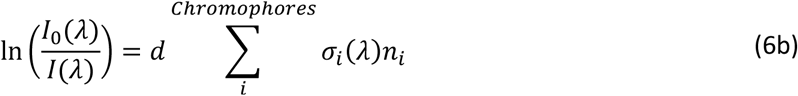

with *d* being the path length of the light through the absorbing volume (see below for units) and *c_i_* the concentration in mol/l. For the single-complex case here, eq. (6b) is more convenient, rescaling the molar (decadic) extinction coefficient to the attenuation cross section *σ* = ln(10) *ϵ/N_A_, N_A_* being Avogadro’s number. The corresponding concentration *n* is then simply the number of chromophore molecules in the considered volume. For CP29, we estimated the volume by the approximate dimensions of the complex (45*40*35 Å^3^ = 63000 Å^3^). To account for a randomized irradiation direction, this volume is then represented by a sphere with a 25 Å radius (volume of about 65450 Å3). Hence, a pathlength *d* of 50 Å was used.

Obviously, the actual intensity difference Δ*I* between *I*_0_ and *I* is the value of interest, resulting in

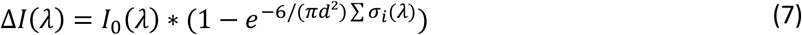

with every chromophore spectrum *σ_i_*(*λ*) shifted according to the CP29 site energies (Table S2 in the SI). For every *λ*, Δ*I*(*λ*) was split linearly according to the contribution of each chromophore to ∑*σ_i_*(*λ*), obtaining the corresponding absorbed intensity per chromophore, Δ*I_i_*(*λ*). Finally, this contribution was integrated over the range between 350 and 550 nm (B band region) to obtain the weight of absorption for each chromophore, relative to the total CP29 absorption:

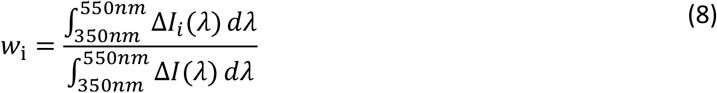

## Results and discussion

### Q and B band spectral overlaps

Computed *J* values for various spectrally unshifted Chl pairings are shown in Table 1. Including the full spectrum is not affecting *J* in most cases, as the values shown in parentheses do not differ much to the original value. This indicates that the potential to directly donate into a different band is minute; even non-existent for Q → B. The strongest B → Q overlap is found for the *b/a* pairing (0.46 10^14^ M^-1^ cm^-1^ nm^4^ arising from including the Q band), indicating a small chance for (Chl *b* B) → (Chl *a* Q) FRET. However, since Chl *b/a* for B-B is only 2.38, compared to up to 27.7 for *a/b*, EET from Chl *b* to *a* can be expected to play an insignificant role.

**Table 1:** J values (in 10^14^ M^-1^ cm^-1^ nm^4^) for unperturbed Chl donor/acceptor pairs, using different spectral ranges (B band only or both B, Q bands).

| Donor/Acceptor | $J_{Q-Q}$ ($J_{Q-(B,Q)}$) | $J_{B-B}$ ($J_{B-(B,Q)}$) |
| --- | --- | --- |
| $a/a$ | 55.1 (55.1) | 12.0 (12.1) |
| $a/b$ | 7.74 (7.74) | 27.6 (27.7) |
| $b/a$ | 41.0 (41.0) | 1.92 (2.38) |
| $b/b$ | 41.7 (41.7) | 24.0 (24.4) |

The results displayed in Table 1 can be easily explained: FRET becomes more efficient at lower energies due to the *λ*^4^ scaling (eq. (5)). Further, Chl *b* absorption shows a stronger maximum than Chl *a* in the region band (ratio 1.42 *b/a* at *λ_max_*). Both effects result in *J* for Chl *b*/*b* homotransfer to be twice as large compared to Chl *a*/*a* (24.0 vs. 12.0). When shifting the Chl *b* absorption and emission spectra in a test calculation to the Chl *a* energies (+0.162 eV and +0.152 eV, respectively), the Chl *b* homotransfer *J* becomes 17.9. This shows that 5.9 of the difference in *J* between the *a/a* and *b*/*b* transfers arises from the spectral shapes, and 6.1 from the energetic shift.

At this point, the nearly-perfect fit between the spectral shapes of the Chl *a* B band emission and the Chl *b* B band absorption should be acknowledged (see Leupold *et al*^41^), which leads to the high *J* of the Chl *a/b* interaction (27.6). Just based on the spectral overlaps, FRET apparently favors EET to Chl *b* above all other B-B processes. Furthermore, once located at Chl *b*, back-transfer to Chl *a* appears to be almost impossible (1.92).

### Q and B band Förster radii

R_0, B-B_ (and R_0, Q-Q_ for comparison) for Chl *a* and *b* pairings can be computed from Eq. (4) using different *κ*^2^, the results being shown in Figure 2 and Table S5 of the SI. The range of EET, expressed as R_0_, depends strongly on the alacrity of the decay process 1/*τ_D_*: For Chl *a* monomers, *τ_D,Q_* in the range of 6 ns,^46,62^ while *τ_D,B_* of Chl *a* is ~100 – 250 fs, for Chl *b* 58 fs.^23–25^ Efficient EET (Eq.(1)) rates (Eq.(2)) must exceed the respective 1/*τ_D_* by one or more orders of magnitude. This then translates to a high value of R_0_.

**Figure 2:**
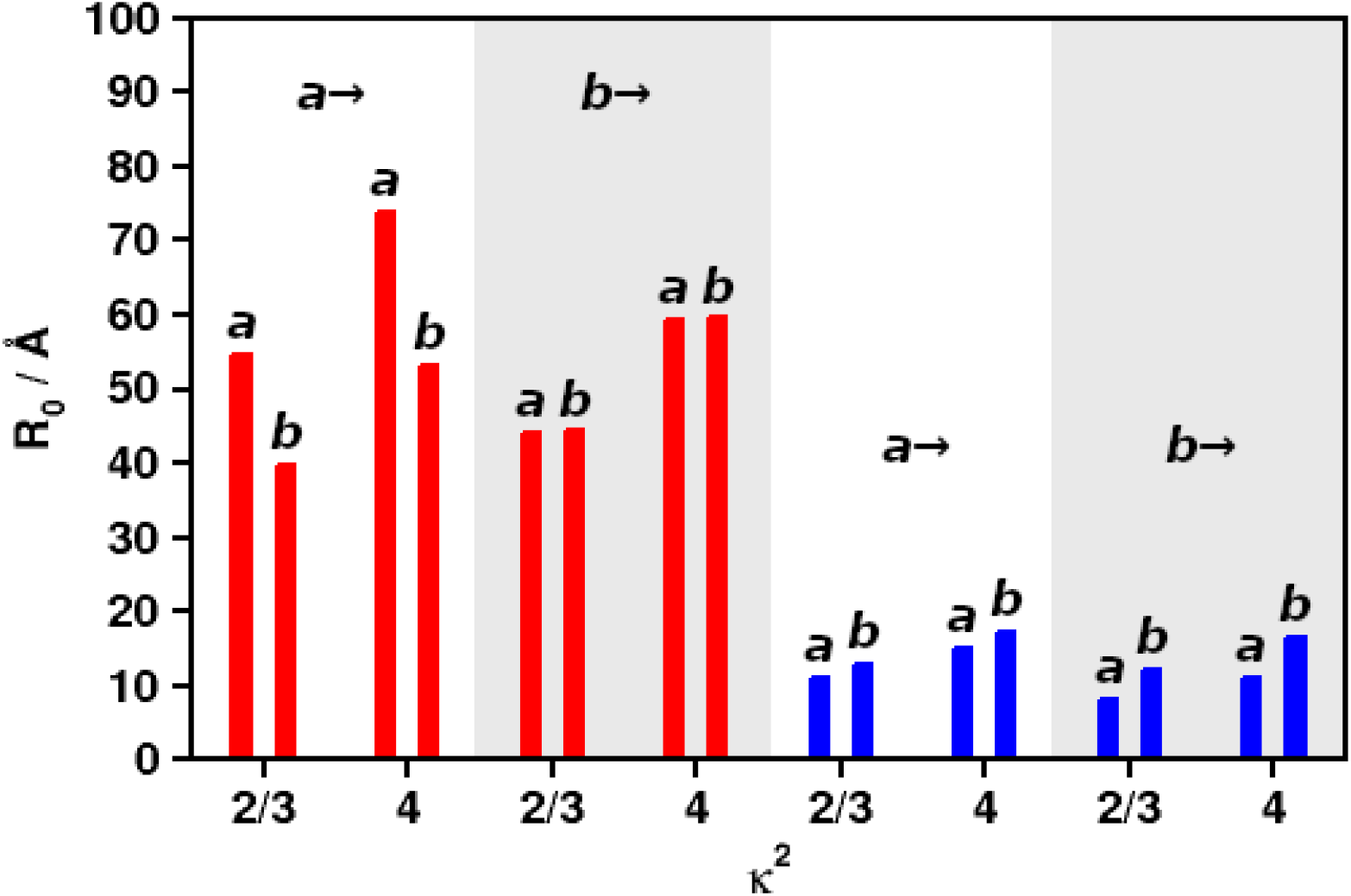
R_0_ for Q-Q (red) and B-B (blue) FRET, using different values of *κ*^2^ and different donor/acceptor combinations of Chls a/b (left/right in each column pair). Donors indicated above each column quartet.

Figure 2 indicates that Chl *a* has the highest priority (R_0, Q-Q_ between 54.7 and 73.8 Å depending on *κ*^2^) as an acceptor for Chl *a* emission in the Q band (red bars, first quartet “*a*→”). For Q band Chl *b* donation, there is virtually no preference (red bars, second quartet “*b*→”). The model predicts the experimental range of Chl *a/a* transfer (55-75 Å^49,63^) and lower-*κ*^2^ Chl *b*/*b* (computed 44.2 vs exp. 46 Å^49^) very well. In contrast to the Q band, B band EET (blue bars) clearly prefers Chl *b* as acceptor in all cases. Figure 2 basically agrees with the picture already seen from the *J* values in Table 1, with the Chl *b* being the preferred acceptor Chl in the B band and Chl *a* in the Q band, at least for Chl *a* as the donor.

As in the first part of this article series, it needs to be pointed out that the small R_0, B-B_ values do not indicate that B-B FRET is actually inefficient. Efficient FRET still occurs if a high number of acceptor molecules is nearby, provided that the *sum* of all *k_FRET,DA_* strongly exceeds 1/*τ_D_* (Eq. (1)). To investigate such a case, the CP29 model will be taken as an example below.

### B band EET network in CP29

In the previous article of the series, a general assessment of EET in the B band region of CP29 was presented. Here, efficiencies for EET to different pigment classes are shown in more detail (Figure 3; for detailed values, see previous article in this series). It can be seen that a large amount of the B-B EET (blue circle segments; dark blue to Chl *a*, light blue to Chl *b*) is actually to or between Chl *b*. Despite only representing 4 out of 14 Chls, Chl *b* is actually the preferred acceptor for B-B EET: For all CP29 Chls, 26.0% of all B band processes shown in Figure 3 (Averages, “CP29”) have Chl *b* as EET acceptor. In contrast, Chl *a* is only the acceptor in 21.3% of all cases, despite being 2.5 times higher in number than Chl *b*. Especially strong donation into Chl *b* show Chls *a* 604 and 613 (93.0% and 90.2%, Figure 3, large circles, center-bottom), Chl 609 and 610 also display significant Chl *a/b* B-B EET (both 21%). Except for Chl *b* 614, which has no CP29 Chl *b* receptors nearby, all Chls *b* nearly exclusively donate B band energy between each other (606: 35%; 607: 84%) or to Crts (606: 51%; 608: 84%).

**Figure 3:**
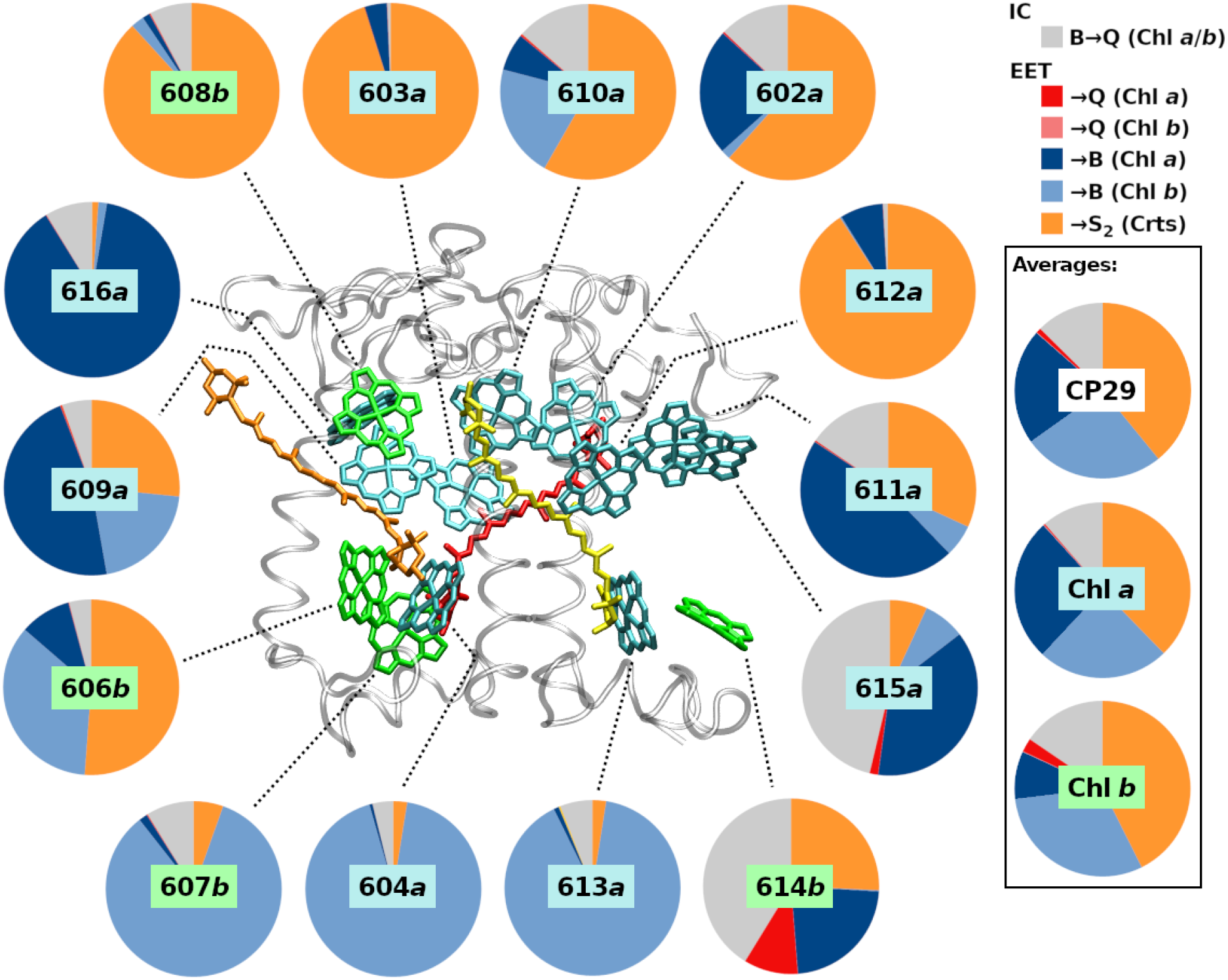
E_FRET,total_ fractions for each B band excited CP29 Chl donor, showing all classes of IC and FRET processes.

The high preference for Chl *b* acceptors is likely due to the remarkable spectral identity of the Chl *b* B band absorption with Chl *a* B band emission mentioned above, as observed in the work by Leupold *et al*.,^41^ leading to the strong *J_B–B_* and larger R_0, B-B_ for the B band (see previous sections). It could however still be that site energies define the role of the bound chromophore as donor/acceptor, which is addressed below.

### Chl exchange and site energy test

To check if site energies result in the B band acceptor role of bound chromophores,, two tests were conducted, (i) replacing all CP29 Chl *b* by Chl *a* and (ii) removing all site energy shifts. The detailed results of test (i) can be found in the SI (Figure S1 for changes in the Q band and S_2_ for changes in the B band). For test (ii), we restrict the discussion here to the averages of the B band for sake of brevity. If the site energies are the determining factor, we would expect Chl *a* to also act as B band acceptors in test (i) and an equalization of EET preference in test (ii).

The first test, Chl exchange, results for the Q band in a reweighting between Q band acceptors; EET efficiency to the Chl *a* Q state sites drops from 78% to 71% in favor of Chl *b* sites (in the Q band, there are no significant other processes, even IC is vastly outpaced by EET). For the B band (Figure S2), it is found that the Chl *b* sites are much less prone to accept B band energy when occupied by Chl *a* (20.0% instead of 26.0% donation into Chl *b* sites vs. 27.8% instead of 21.3% donation into Chl *a* sites). For the Chl 606*b* site, even a strong donation to the Chl 604*a* site is found (up from 9% to 39%). Note that Figure S2 shows some drastic changes for the Chl 614*b* site upon Chl exchange; this however is to be taken with care as Chls 614 and 615 are likely coupled to pigments that are absent in the CP29-only model (*e.g.*, pigments from the neighboring CP24 complex).

Removing the site energy shifts, test (ii), results in a slight drop in Chl *b* B band acceptor role (24.8% instead of 26%; Chl *a*: 20.8% without instead of 21.3% with site shifts). The effect on Chl *b* sites is thus much smaller (1.2%) than the effect Chl exchange (6.0%). This shows that the Chl *b* B acceptor preference originates mostly from the pigment type and only weakly from the site energy. In the light of the strong *a/b J_B–B_*, this is not surprising.

It is thus already possible to conclude here that Chls *b* are preferred acceptors of B band energy, also in CP29, despite their lower numbers. Further, once located at Chl *b*, B band energy for the most part has to either stay at Chls *b* or to transfer to Crts. Functionally, Chl *b* traps B band energy in the peripheral antennae, as Chl *b* does not occur in the RCs and B-B EET to Chl *a* is energetically uphill.

### Spectral competition for initial B band absorption

The results from the previous article in this series, together with the above insights, yield the conclusion that Crts and Chl *b* suppress Chl *a* B-B EET. The sequence of events upon CP29 illumination can be stated as follows: (i) Spectral competition between Crts, Chls *b* and and Chls *a*, (iia) EET from Chl *a* to Chls *b* and Crts, (iib) EET from Chl *b* to Crts and finally (iii) Q band EET from Crts to Chl *a* (EET from Crts to Chl *b* Q is only 5% for Vio, 4% for Neo and 1% for Lut; data not shown; this process will therefore not be discussed).

The first step in the sequence, spectral competition, has so far not been considered in detail here. It can be evaluated using Eq. (8), with the results listed in Table S6 of the SI. A graphical representation of the results for terrestrial sunlight irradiation^20^ is given in Figure 4 (left circles in each subgraph, “Abs.”). Direct Chl *a* absorption only makes up for 39.6% of the total absorption probability for full CP29 (dark blue in Figure 4, top left subgraph). Using white light (see SI) increases this fraction to 47.7%. Despite only 4:10 Chl *b/a* pigments in CP29, Chl *b* absorption is 25.4% in sunlight and 22.0% in white light. This indicates that Chl *b* is able to efficiently compete with Chl *a* absorption, especially under regular sunlight conditions.

**Figure 4:**
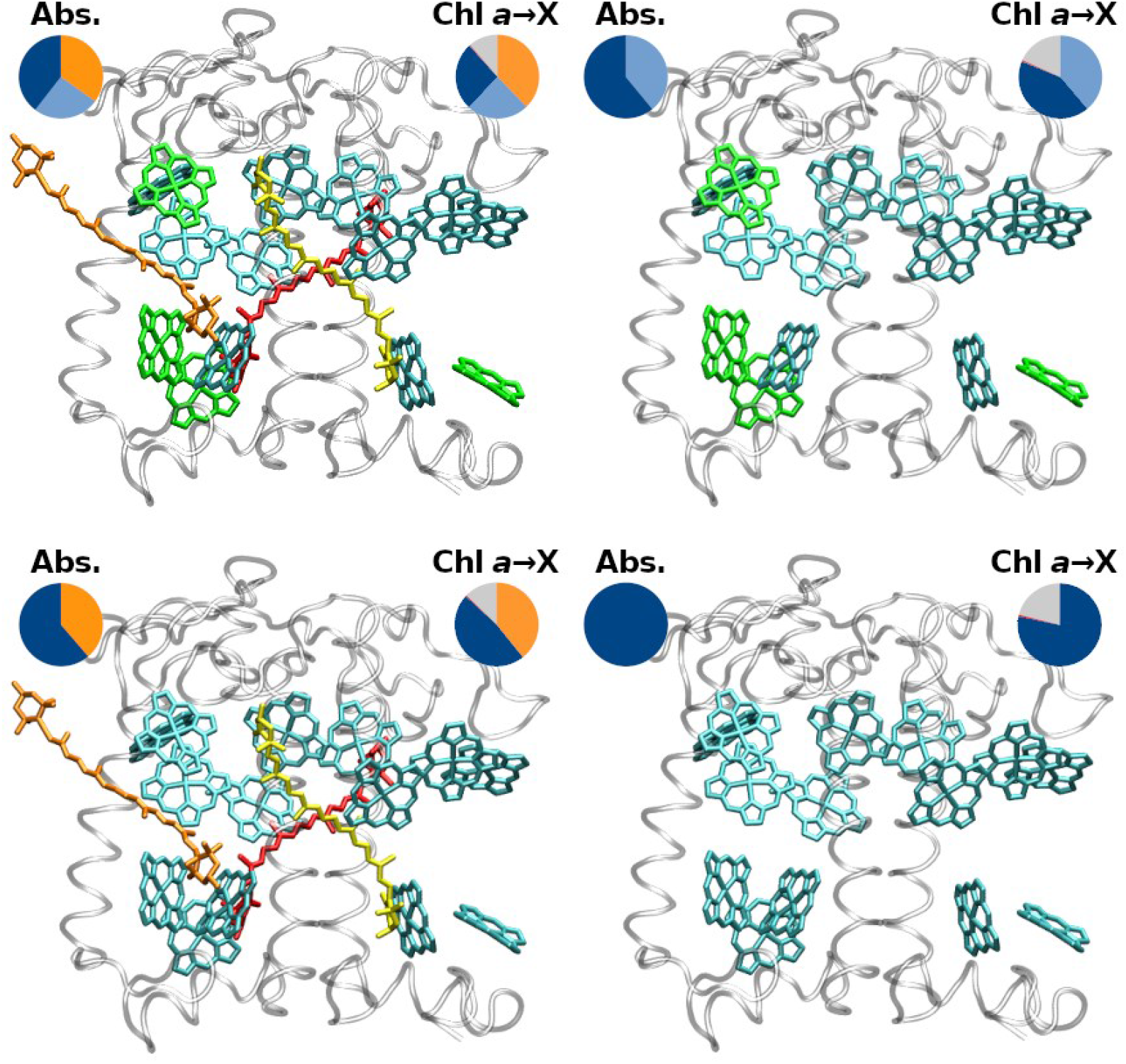
Tested CP29 pigment combinations and resulting absorption ratios for terrestrial sunlight (left circles) and averaged Chl a B band IC/EET event ratios (right circles). Chl absorption colors analogous to IC/EET colors as in Figure 3. Top row: WT CP29 (left) and Crt-free CP29 (right); bottom row: Chl b replaced by Chl a, with Crts (left) and without Crts (right).

For terrestrial sunlight, as shown in Figure 4, removal of the Crts (top right) yields a Chl *a* absorption fraction of 60.9%, while replacement of Chl *b* by Chl *a* (bottom left subgraph) in the presence of Crts changes this fraction to 61.3%. The latter change (39.6 to 61.3%) is larger than expected, considering that 4 Chls *a* are effectively added, increasing its concentration by only 40%. However, the site energies of some Chl *b* sites, especially 606 and 607 (see Table S2 in the SI) are not well covered by the original Chl *a* shifts. The newly added Chl *a* therefore compete much less with other chromophores in the system, leading to the comparatively high gain in absorption weight.

Our absorption analysis has direct consequences for any experiment exciting in specific regions of the B band. The experiments by Leupold and coworkers^41^ offer a unique perspective due to the indirect excitation. Here, no interference from Crts can be expected, only Chl/Chl competition in the Q region, which is screened out by the selective excitation from Q to B. However, it is difficult to assess the results without a simulation of the LHCII complex, which they compared to CP29. Corresponding calculations are currently under way.

Another example for blue light interaction within CP29 are the experiments of Crimi *et al*.^64^ They used excitation wavelengths of 440/475 nm, and reportedly observed no differences in CP29 Q band fluorescence. The two pulses are supposed to preferentially excite Chls *a/b*, respectively, and should yield the distribution of CP29 excitation as shown in Table S7 of the SI. For the Chl *a*-preferring pulse (440 nm), Chl *b* absorption is nearly on par with Chl *a* (*w_Chl a_/w_Chl b_* 35.5%/30.1% for a pulse width of 250 cm^-1^, 37.3%/29.1% for 500 cm^-1^). For the Chl *b*-pulse (475 nm), Chl *b* excitation strongly exceeds that of Chl *a* (*w_Chl a_/w_chl b_* 5.8%/32.9% for 250 cm^-1^, 11.1%/34.4% for 500 cm^-1^), but most excitation is found at the Crts (62.3% or 54.5%) regardless of the pulse widths. Summed up, the main excitation is either found in the Chls (440 nm pulse, about 66%) or the Crts (475 nm pulse). The observed indifference for the Q band fluorescence measured by the experiment can thus only be explained by our proposed B band EET pathways, as localized IC in Crts and Chls should yield significant differences. Within our model of “assisted” IC, the original site of excitation is not of high relevance, as nearly all excitation gets funneled via Chl *b* or Crts at some point (see also Figure 5 below).

**Figure 5:**
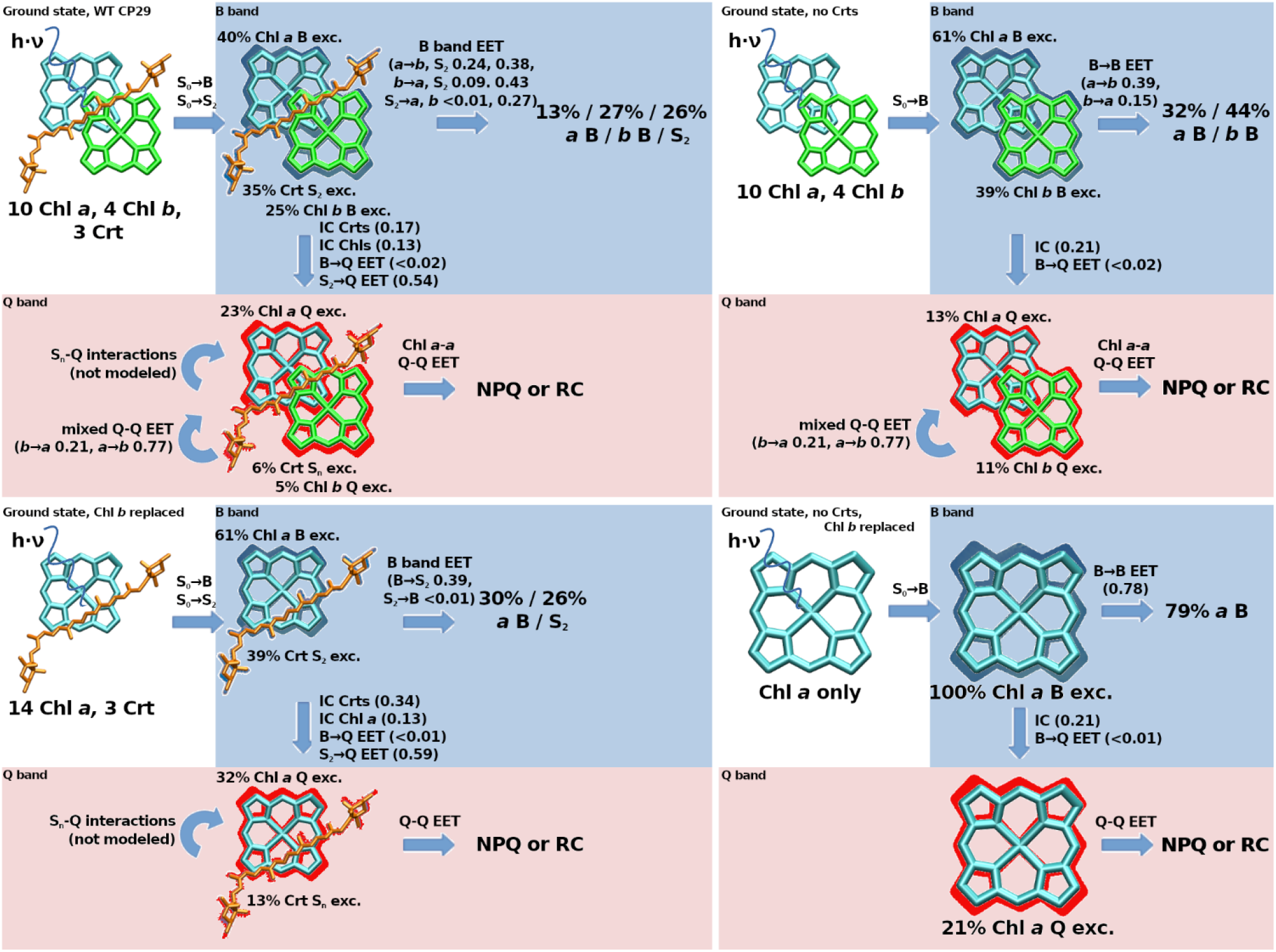
Computed populations resulting from the initial absorption step in terrestrial sunlight (Figure 1A) and subsequent FRET redistribution in different versions of the CP29 complex’ chromophore compositions. Same systems/order as in Figure 4. S_n_ refers to any state other than S_2_.

### Combined effect of the accessory pigments on the B band

The analyses might indicate that only 39.6% of all B band excitations affect Chl *a* in the first place, which would make the presented EET protection pathways by Crts (cf. previous article) and Chl *b* (see above) much less necessary. Still, in the FRET model, these 39.6% Chl *a* population will be reduced by a factor of 0.66 for each IC/EET step (0.24 from EET to Chl *b*, 0.38 from EET to Crts and 0.11 from IC, see right circle in top left graph of Figure 4, slightly offset by repopulation mainly from Chl *b*, resulting in 0.66 effective loss). This means that on average, each EET step only retains a fraction of 0.34 of the initial Chl *a* B state population, which would eliminate 99% of any initial B band population after about 5 “jumps”. As can be seen, all other tested pigment combinations in Figure 4 perform much worse, only removing about half of the Chl *a* B band population for each IC/EET event (top right and bottom left graphs in Figure 4, right circles). A CP29 containing only Chl *a* would have to rely on IC for B band removal, which is outpaced by EET by about 3:1 EET:IC (bottom right graph of Figure 4, right circle). For such a Chl *a*-only system, this would mean that 99% removal is only ensured after 26-27 “jumps” (disregarding the fact that such a system has no choice other than Chl *a* as initially excited chromophore, foregoing on the initial B band competition effect). These processes are shown in detail in Figure 5 for the absorption and initial redistribution steps.

Figure 5 indicates a balanced view on the roles of Crts and Chl *b*. Crts alone would keep a large proportion (13%, bottom left graph, Q band) of excitations without EET back to the Q band. It may be that other Crt states, such as the dark S_1_ state - not covered in the model - will provide additional back-transfer.^65^ However, within the model, Chl *b* provides the possibility for Crts to donate back to the Chl *b* B band, leading to 27% B band population (top left graph, after B band EET). As a consequence, the Crt population in the Q band drops drastically, now 6%, compared to 23% Chl *a*. Without Chl *b*, the Q band Chl *a*:Crt ratio is 32%:13%, including Chl *b* should thus allow for an improved quantum yield from B band excitations. These effects are mainly associated to the Chl *b*-tuned emission spectrum of Neo; it will be interesting to see in future models how this differs for Neo-free systems. The right-hand side of Figure 5 indicates that the presence of Chl *b* will slightly increase B band depletion (79%/73% remaining population with/without Chl *b*), which is due to its slightly larger decay rate (B → Q IC). Its primary function on its own, however, appears to be the initial absorption competition, with the result of trapping 2/3 of the Chl B band excitations in the peripheral antennae (13% vs 27%, top left graph after EET).

## Conclusions

An assessment of the B band absorption and EET events in CP29 has been presented. While the previous article in this series focused on the effects of Crts on B band EET, this article mainly concerns Chl *b* and the initial absorption competition.

It can be concluded that the accessory pigments in CP29 prevent Chl *a* B band excitation very efficiently, even at the initial absorption stages: under sunlight conditions, only 20% of initial B band region excitations would be located at the Chls *a*. The model assumes almost no reverse EET to Chls *a*, which is likely an exaggeration due to the instant pigment reorganization in the model. Within this framework, most Chl *a* B band excitations are relocated to Crts or Chls *b* via EET. B band EET larger than 10% to other Chls *a* is only found for Chls 602, 609, 611, 615 and 616. Including additional PSII supercomplex pigment-proteins will likely reduce the B-B EET even for those pigments, as was shown in the previous article of this series. Associated complementary calculations are running in our lab. Taking only CP29 within the present FRET model, the 40% Chl *a* population will be reduced by a factor of 0.66 by each IC/EET step.

It must, however, be noted that EET back to the Chls *a* is still possible, even if the overlap was found to be small (see Table 1). The instant reorganization to the emission spectrum may be a gross oversimplification given the small timescales involved. It is likely that immediately after EET to Crts and Chl *b*, the corresponding overlap for back-donation to Chl *a* absorption is higher. As such, the actual B state population of Chls *a* can be expected to be much more dynamic, as transfer to Chl *a* will likely occur to a higher degree than in the presented model. Also, formation of excitons for coherent EET has not been covered in our model.

It is beyond the scope of this article to assess how much B band excitation can actually be tolerated by the RCs. Given the almost overbearing protection that the accessory pigments provide in terms of the B band, it would suggest that even trace amounts of B excitation could be a problem. From an evolutionary perspective, this is easily understandable – in early, less crowded pigment-protein complexes with larger intermolecular distances, only Q band EET is efficient; B → B EET likely is a problem that occurred later, when complexes approached a higher EET efficiency threshold.

It would be beneficial for an organism if the protection described here would also be valid for not perfectly assembled PSII variants. Our test calculations indicate that B band protection would still be intact if individual Crts or Chls *b* were removed/exchanged (data not shown; will be covered in a future article). The total effect of B band protection is thus not due to individual chromophores, but due to many factors, making the mechanism extremely robust against mutations or assembly errors.

What is obviously missing here is the actual amount of B band excitations flowing out of CP29 to other complexes, most importantly the RCs. It is very important to ask how much protection the RC itself provides in terms of B band absorption. The RCs only contain Chls *α* and β-carotene, and the absence of Chl *b* is necessary – when assuming the role of a B band trap. Note that an earlier article indicated that the protein environment is relevant in this aspect, too (see also Table S8 in the SI, showing that state-specific energy shifts are not trivial phenomena).^43^ Possibly, without the help of accessory pigments, nature had to rely on a protein based EET steering even more in the RCs than for CP29.

All these important questions have to be answered in future research. It should, however, be noted that already the appearance of EET between higher states may indicate a possible use as an untapped energy source for biotechnological applications.

## Supporting information

Supplementary material (all)

## Acknowledgements

J.P.G. gratefully acknowledges funding by the Deutsche Forschungsgemeinschaft (German Research Foundation, DFG), grant number 393271229. H.L. gratefully acknowledges funding by the Czech Science Foundation, GAČR (grant no. 22-17333S). The authors thank Prof. Dr. Peter Saalfrank for his valuable comments on the manuscript.

## Author contributions

J.P.G. provided the theoretical concept, calculations, and figures, as well as the initial draft. H.L. provided conceptual guidance, the experimental spectra and edited the draft. Both authors contributed equally to the finalization of the manuscript.

