## Supplementary material (all) for "Excitation energy transfer between higher excited states of photosynthetic pigments: 2. Chlorophyll *b* is a B band energy trap"

**Supporting Information**

Jan P. Götze[1,*] and Heiko Lokstein[2]

*[1]Institut für Chemie und Biochemie, Freie Universität Berlin, Arnimallee 22, 14195 Berlin, Germany; ORCID ID 0000-0003-2211-2057*

*[2]Department of Chemical Physics and Optics, Charles University, Ke Karlovu 3, 121 16 Prague, Czech Republic; ORCID ID 0000-0001-6739-4612*

Table of contents:

- Zn-porphyin Soret EET
- Pigment fluorescence yields
- CP29 site energies
- CP29 structure distance matrix
- CP29 Q band EET changes upon Chl *b* replacement
- CP29 B band EET changes upon Chl *b* replacement
- CP29 B band absorption probabilities per pigment
- CP29 excitation patterns after laser excitation
- Chl *a* maxima of Q and B bands shift similarly in different environments

*Zn porphyrin Soret EET*

Work on Zn-porphyrin windmill-like arranged covalently connected structures is instructive.[1] The authors attribute their findings of possible Soret-Soret EET to the special properties arising from Zn coordination. They also state the prerequisites for efficient EET, namely "… it has the close proximity of the donor and acceptor that is a prerequisite to [Soret-Soret] energy transfer, as well the very favorably matched spectral overlap between the fluorescence emission from the [Soret] state…". They analyze the splitting between the coupled Soret states and correlate it to the orientation of the involved transition dipole moments (TDMs) using a FRET model. Nakano *et al.* [2] investigated also covalently bound, benzene-bridged porphyrine trimers and basically arrive at the same conclusion as for the windmill-like arrangement.

Karolczak *et al.* [3] characterize the Soret emission of Zn porphyrins further. It is found that the Stokes shift is very small (115 $cm^{-1}$) and the emission is relatively strong for a non-$S_1$ emission, with a quantum yield (in ethanol) of $1.84 \times 10^{-3}$.

*Pigment fluorescence yields*

*Table S1: Fluorescence yields of Chl B bands and Crt $S_2$ used for the calculations. References see main text.*

| | Chl *a* | Chl *b* | Lut | Vio | Neo |
|---|---|---|---|---|---|
| B band yield | 1 $10^{-4}$ | 0.92 $10^{-4}$ | 1.5 $10^{-4}$ | 1.5 $10^{-4}$ | 1.5 $10^{-4}$ |

*CP29 site energies*

*Table S2: $Q_y$ and $B_X$ energy shifts in CP29 derived from data in Petry and Götze, relative to averages of Chl a or b states/sites.*

| **Site ID** | **602a** | **603a** | **604a** | **606b** | **607b** | **608b** | **609a** |
|---|---|---|---|---|---|---|---|
| **Q shift / meV** | -11 | +1 | +7 | +17 | -8 | +8 | +21 |
| **B shift / meV** | -32 | +22 | -1 | +40 | +55 | -54 | +19 |
| **Site ID** | **610a** | **611a** | **612a** | **613a** | **614b** | **615a** | **616a** |
| **Q shift / meV** | +3 | -24 | +18 | -1 | -17 | -51 | +40 |
| **B shift / meV** | -4 | -65 | +15 | +32 | -43 | -76 | +93 |

*CP29 structure distance matrix*

*Table S3: CP29 distance matrix, according to chain R of PDB entry 5XNL, Mg-Mg distances, in Angstrom*

| **603** | **604** | **606** | **607** | **608** | **609** | **610** | **611** | **612** | **613** | **614** | **615** | **616** | |
|---|---|---|---|---|---|---|---|---|---|---|---|---|---|
| 11.7 | 24.9 | 26.8 | 24.9 | 21.5 | 17.9 | 15.9 | 15.6 | 17.9 | 18.9 | 24.4 | 13.1 | 20.7 | **602** |
| | 19.3 | 18.3 | 15.1 | 17.8 | 9.3 | 18.3 | 24.5 | 22.8 | 19.6 | 28.1 | 22.9 | 13.0 | **603** |
| | | 7.9 | 11.7 | 18.2 | 17.4 | 18.3 | 26.8 | 19.2 | 20.6 | 26.6 | 32.0 | 27.0 | **604** |
| | | | 9.5 | 15.9 | 13.7 | 20.8 | 32.2 | 25.2 | 25.8 | 33.1 | 36.1 | 22.5 | **606** |
| | | | | 22.0 | 15.3 | 24.8 | 32.0 | 26.9 | 20.8 | 29.2 | 33.0 | 22.3 | **607** |
| | | | | | 10.4 | 11.5 | 27.9 | 22.2 | 30.8 | 37.2 | 33.1 | 17.2 | **608** |
| | | | | | | 16.5 | 28.6 | 24.6 | 26.2 | 34.3 | 30.1 | 9.82 | **609** |
| | | | | | | | 16.9 | 11.7 | 24.6 | 29.0 | 24.4 | 23.7 | **610** |
| | | | | | | | | 9.2 | 18.6 | 18.0 | 12.3 | 34.5 | **611** |
| | | | | | | | | | 18.3 | 19.4 | 19.7 | 32.4 | **612** |
| | | | | | | | | | | 9.2 | 17.3 | 32.6 | **613** |
| | | | | | | | | | | | 18.1 | 40.9 | **614** |
| | | | | | | | | | | | | 33.4 | **615** |

*Table S4: CP29 Chl-Crt distance matrix, according to chain R of PDB entry 5XNL, Mg-Crt(COM) distances, in Å. Crt(COM) (center of mass) computed for conjugated carbon atoms present for each Crt in 5XNL.*

| | **602** | **603** | **604** | **606** | **607** | **608** | **609** | **610** | **611** | **612** | **613** | **614** | **615** | **616** |
|---|---|---|---|---|---|---|---|---|---|---|---|---|---|---|
| **Lut** | 16.7 | 18.1 | 13.0 | 18.6 | 20.6 | 17.7 | 19.1 | 9.7 | 14.2 | 6.6 | 16.6 | 20.5 | 21.8 | 27.7 |
| **Neo** | 29.2 | 23.8 | 14.9 | 12.2 | 21.3 | 10.0 | 16.3 | 17.3 | 32.8 | 25.1 | 33.2 | 39.2 | 39.5 | 24.4 |
| **Zea** | 11.2 | 9.6 | 15.7 | 18.8 | 15.5 | 20.6 | 15.6 | 16.3 | 17.6 | 15.6 | 11.0 | 18.9 | 17.7 | 22.2 |

*Spectral parameters of Chl-Chl FRET*

*Table S5: FRET parameters computed for Q-Q and B-B EET in Chls a and b. $J$ in $10^{14}$ $M^{-1}$ $cm^{-1}$ $nm^4$, $R_0$ in Å.*

| **Homotransfer** | $J_{Chl\ a/Chl\ a}$ | $J_{Chl\ b/Chl\ b}$ | $R_{0,Chl\ a/Chl\ a}$ | | $R_{0,Chl\ b/Chl\ b}$ | |
|---|---|---|---|---|---|---|
| $\kappa^2$ | / | / | 2/3 | 4 | 2/3 | 4 |
| **Q band** | 55.1 | 41.7 | 54.7 | 73.8 | 44.2 | 59.6 |
| **B band** | 12.0 | 24.0 | 11.1 | 14.9 | 12.2 | 16.5 |
| **Heterotransfer** | $J_{Chl\ a/Chl\ b}$ | $J_{Chl\ a/Chl\ b}$ | $R_{0,Chl\ a/Chl\ b}$ | | $R_{0,Chl\ a/Chl\ b}$ | |
| $\kappa^2$ | / | / | 2/3 | 4 | 2/3 | 4 |
| **Q band** | 7.74 | 41.0 | 39.5 | 53.2 | 44.1 | 59.4 |
| **B band** | 27.6 | 1.92 | 12.7 | 17.1 | 8.8 | 10.8 |

*CP29 Q band EET changes upon Chl b replacement*

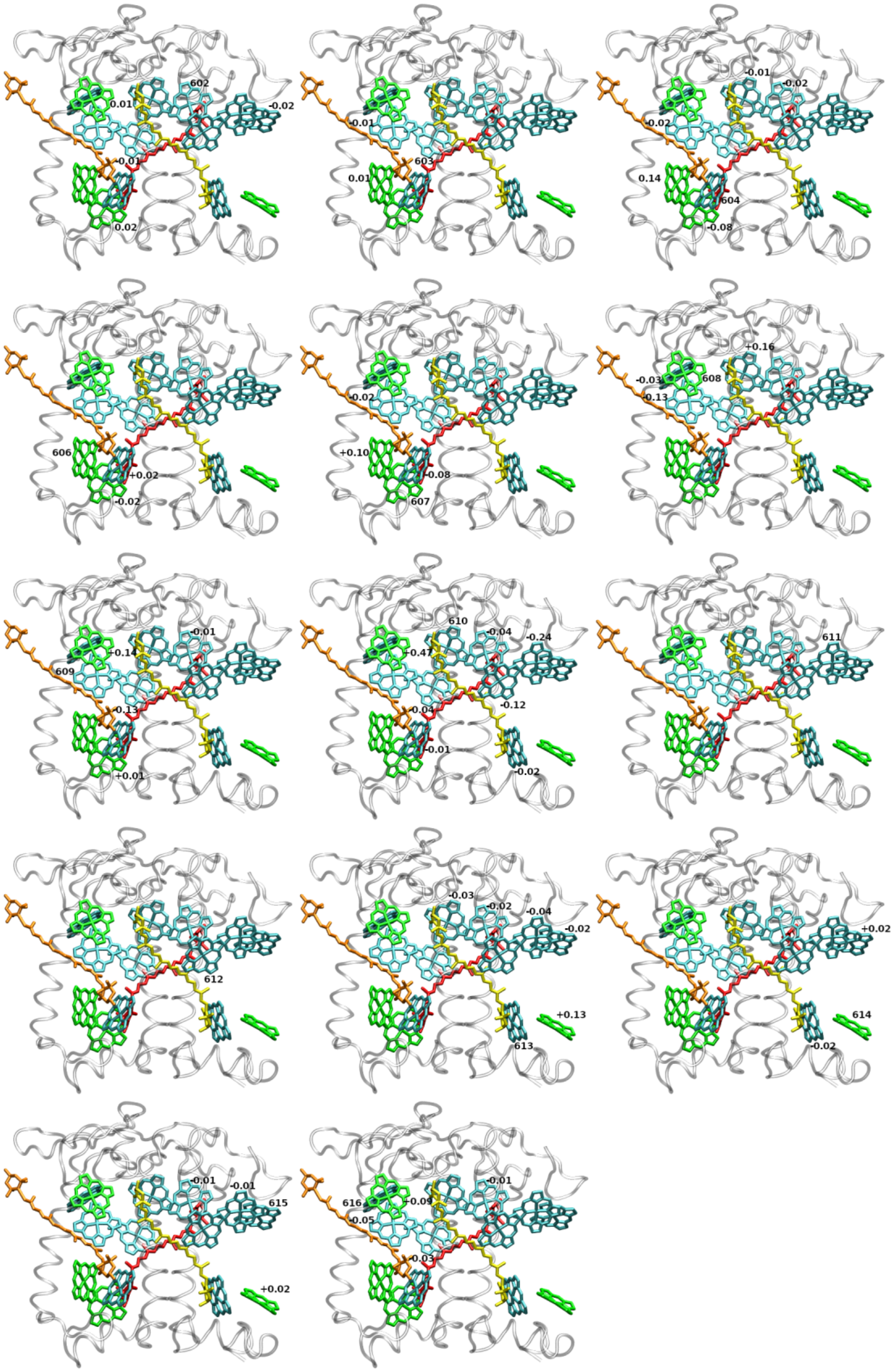


*Figure S1: Differences in CP29 FRET efficiencies for Chl B band donation to all possible acceptor bands. Current Chl index shown. Explicit individual EET changes for each acceptor, depicted for each donor Chl in the system. Pigments shown for the wildtype chromophore configuration (Chl b present).*

*CP29 B band EET changes upon Chl b replacement*

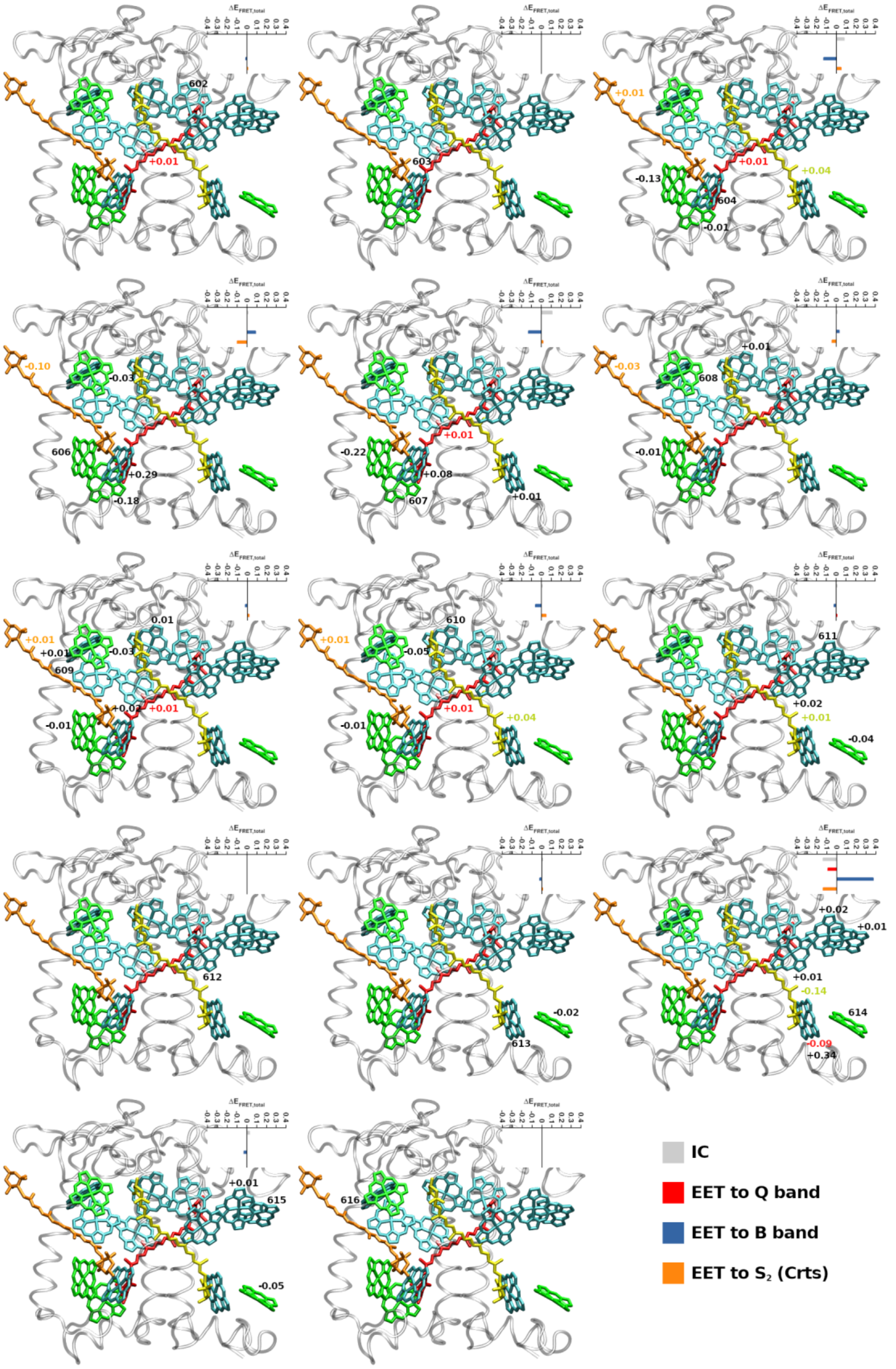


*Figure S2: Differences in CP29 FRET efficiencies for Chl B band donation to all possible acceptor bands. Small graphs: summed differences for each EET class (see legend in lower right corner). Current Chl index shown in structural model. Structural models: Explicit individual EET changes for each acceptor, depicted for each donor Chl in the system. Pigments shown for the wildtype chromophore configuration (Chl b present).*

*CP29 B band absorption probabilities per pigment*

*Table S6: Relative weights of chromophore groups in B band CP29 absorption. Integration from 350 to 550 nm, with B band Chl absorption as given in the main text. W: White light (equal intensities over all wavelengths), S: terrestrial sunlight intensity distribution as shown in Figure 2A of the main article, with maximum I = 1 at 528 nm.*

| | **Chl *a*** | **Chl *b*** | **Lut** | **Vio** | **Neo** |
|---|---|---|---|---|---|
| $n_i$ | 10 | 4 | 1 | 1 | 1 |
| **W** | 0.477 | 0.220 | 0.110 | 0.093 | 0.100 |
| **S** | 0.396 | 0.254 | 0.132 | 0.109 | 0.109 |
| $n_i$ | 10 | 4 | / | / | / |
| **W** | 0.684 | 0.316 | / | / | / |
| **S** | 0.609 | 0.391 | / | / | / |
| $n_i$ | 14 | / | 1 | 1 | 1 |
| **W** | 0.687 | / | 0.114 | 0.096 | 0.103 |
| **S** | 0.613 | / | 0.146 | 0.120 | 0.121 |

*CP29 excitation patterns after laser excitation*

*Table S7: Relative weights of chromophore groups in B band wildtype CP29 absorption. Integration from 350 to 550 nm, with B band absorption spectra as stated in the main text. Irradiation from a laser pulse focused on $\lambda_0$, with a Lorentzian broadening characterized by the full width half maximum (FWHM).*

| **$\lambda_0$/nm** | **FWHM/cm$^{-1}$** | **Chl *a*** | **Chl *b*** | **Lut** | **Vio** | **Neo** |
|---|---|---|---|---|---|---|
| 440 nm | 250 | 0.355 | 0.301 | 0.119 | 0.113 | 0.110 |
| | 500 | 0.373 | 0.291 | 0.119 | 0.109 | 0.108 |
| 475 nm | 250 | 0.058 | 0.329 | 0.248 | 0.201 | 0.165 |
| | 500 | 0.111 | 0.344 | 0.221 | 0.174 | 0.150 |

*Chl a maxima of Q and B bands shift similarly in different environments*

*Table S8: Band maxima of Chl a in different solvents. Shifts are directly correlated to solvent refractive index n; state-specific changes are not observed. Measurements at RT, for methods see Götze (2022).*

| Solvent | $n$ | Band maxima / cm$^{-1}$ | | $\Delta E$(diethylether) / cm$^{-1}$ | |
|---|---|---|---|---|---|
| | | Q | Soret | Q | Soret |
| Diethylether | 1.353 | 15105.7 | 23282.9 | 0 | 0 |
| Hexane | 1.375 | 15128.6 | 23310.0 | 22.9 | 17.1 |
| Acetone | 1.3588 | 15105.7 | 23255.8 | 0 | 27.1 |
| DMSO | 1.479 | 15015.0 | 23041.5 | 95.7 | 241.4 |
| Chinolin | 1.625 | 14914.2 | 22831.0 | 191.5 | 451.9 |
